# The channel activities of the full-length prion and truncated proteins

**DOI:** 10.1101/2022.01.12.475988

**Authors:** Jinming Wu, Asvin Lakkaraju, Adriano Aguzzi, Jinghui Luo

## Abstract

Prion disease is a fatal neurodegenerative disorder, in which the cellular prion protein PrP^C^ is converted to a misfolded prion which in turn is hypothesized to permeabilize cellular membranes. The pathways leading to toxicity in prion disease are not yet completely elucidated and whether it also includes formation of membrane pores remains to be answered. Prion protein consists of two domains: a globular domain (GD) and a flexible N-terminus (FT) domain. Although a proximal nine polybasic amino acid (FT(23-31)) sequence of FT is a prerequisite for cellular membrane permeabilization, other functional domain regions may influence FT(23-31) and its permeabilization. By using single-channel electrical recordings, we reveal that FT(23-50) dominates the membrane permeabilization within the full-length mouse PrP (mPrP(23-230)). The other domain of FT(51-110) or C-terminal domain down-regulates the channel activity of FT(23-50) and the full-length mouse PrP (mPrP(23-230)). The addition of prion mimetic antibody, POM1 significantly enhances mPrP(23-230) membrane permeabilization, whereas POM1-Y104A, a POM1 mutant that binds to PrP but cannot elicit toxicity has negligible effect on membrane permeabilization. Additionally, anti-N-terminal antibody POM2 or Cu^2+^ stabilizes FT domain, thus provoking FT(23-110) channel activity. Furthermore, our setup provides a more direct method without an external fused protein to study the channel activity of truncated PrP in the lipid membranes. We therefore hypothesize that the primary N-terminal residues are essential for membranes permeabilization and other functional segments play a vital role to modulate the pathological effects of PrP-medicated neurotoxicity. This may yield essential insights into molecular mechanisms of prion neurotoxicity to cellular membranes in prion disease.

## 1 Introduction

Prion diseases are a group of fatal neurodegenerative disorders characterized by the progressive loss of neurons, including human Creutzfeldt-Jakob disease (CJD), scrapie in sheep and mad cow disease. These diseases are associated with two prion isoforms consisting of 209 amino acids: the physiological/cellular prion proteins (PrP^C^) with mainly α-helical structure and insoluble scrapie isoform (PrP^Sc^) with β-sheet rich structure (1). Though the conformational transition and structural misfolding from soluble PrP^C^ to infectious PrP^Sc^ are assumed to characterize prion propagation, the toxic pathogenic mechanism remains to be explored. It has been shown that aggregated PrP^sc^ fibril alone is not on the proximate cause of the pathogenesis (2,3), and membrane-bound PrP^C^ is essential for prion-mediated neurodegeneration (3,4). Soluble low-molecular weight PrP oligomers are much more infectious and neurotoxic than insoluble PrP^Sc^ fibrils *in vitro* and *in vivo* (5,6). This supports the common mechanism ‘toxic oligomer hypothesis’ (7) for neurodegeneration where the oligomers emerge as the most neurotoxic species (8) through permeabilizing cellular membranes and causing cell dysfunction (9,10).

Single-channel electrical recordings revealed that the membrane poration may be one of the mechanisms for neurotoxicity by prions (11-14). The prion protein consists of a flexible N-terminal domain (FT) and a globular C-terminal domain (GD) (**Fig. 1**). The FT region, an intrinsic disordered domain, plays a crucial role in the pathogenesis of prion diseases (15). The primary nine polybasic amino acids (23-31) have been reported as a prerequisite for channel activity of the truncated PrP protein (ΔCR PrP with 105-125 amino acids deleted) (11,12). It remains to be explored how GD and other FT regions modulate channel activity of the primary N-terminal region. As one functional segment of FT, the octa-repeats (OR) region (residues 51-90) binds to metal ions such as Cu^2+^ or Zn^2+^ (16,17) and influences the domain interaction between FT and GD (18-20). A few other functional segments of FT, like two charged clusters (CC1 and CC2), and a hydrophobic domain (HD), may plausibly regulate channel activity of the primary N-terminal residues. How the different functional segments on FT and GD regulate the channel activity may provide insight into the toxic mechanism of prions.

**Figure 1.**
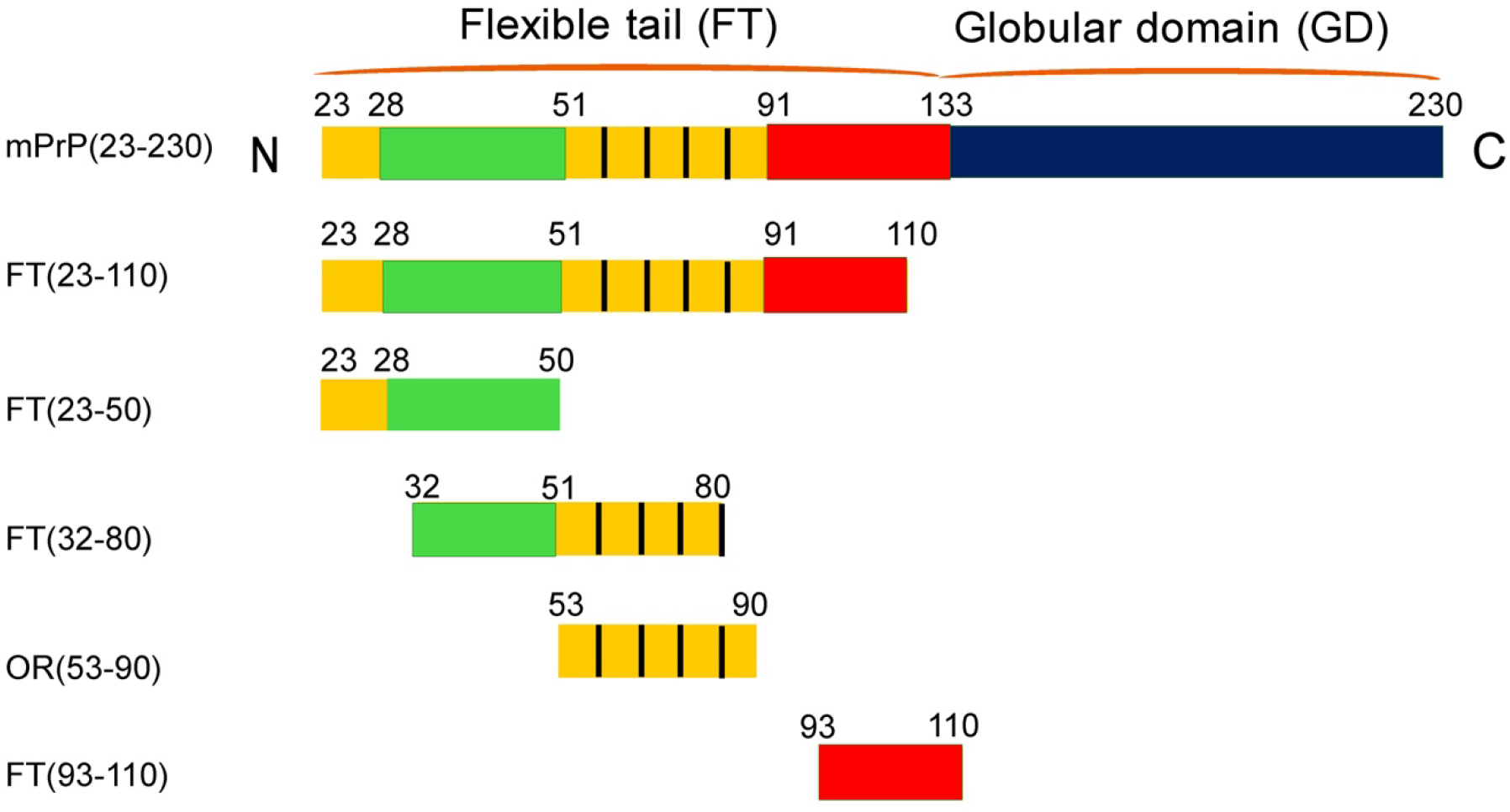
Schematic representation of recombinant mouse full-length mPrP(23-230), consisting of a flexible tail (FT) and a globular domain (GD). The charged clusters CC1 (residues 23-28), CC2 (residues 100-109), octa-repeat region (OR, residues 51-91) and a hydrophobic domain (HD, residues 112-133) are located in the FT domain. PrP fragments, as indicated, were selected to investigate their effects on mPrP ion channel formation in the lipid bilayers.

In this study, we used single-channel electrical recordings to investigate whether the N terminus of the prion protein can induce pore formation and how the different functional segments of FT and GD change channel activity of the primary region. We found that the primary N-terminal domain can form large pores in lipid membranes. In addition, the less channel activity was recorded with a longer sequence starting from the primary N-terminal domain. Further, copper ions and anti-prion antibodies (POM1 or POM2) can stabilize the GD or FT domains and then provoke membranes permeabilization. We therefore reveal that the primary N-terminal domain (FT(23-50)) is essential for permeabilizing cell membranes but other functional segments down-regulate the activity within the full-length mouse PrP (mPrP(23-230)) or other fragments, thus might modulate the pathological effects of PrP-medicated neurotoxicity.

## 2 Materials and methods

### 2.1 Protein expression and purification

The recombinant full-length mouse mPrP(21-230), FT(23-119) and GD(111-231) have been previously described (21). A series of fragmented PrP sequences, including the N-terminal flexible tail FT peptides (FT(23-50), FT(32-80), FT(53-90), FT(93-110)) were all purchased from EZ Biosciences.

### 2.2 Preparation of black bilayer lipid membrane

First, a mixture of DOPC (1,2-Dioleoyl-sn-glycero-3-phosphocholine): DOPG (1,2-Dioleoyl-sn-glycero-3-phospho-rac-(1-glycerol) (20 mg/mL) at a ratio of 4:1 (w/w) which was dissolved in chloroform was prepared in a glass vial and then dried by N_2_ in a fume hood. To thoroughly eliminate the chloroform, a vacuum desiccator was further used overnight. The thin lipid film was added pentane (99%, Sigma-Aldrich, Cat number: 236705) to a final concentration of 10 mg/mL. The lipid bilayer was formed followed by the “Montal-Muller” technique (22). Briefly, a home-made chamber separated by a 25 μm thick Teflon film (Good Fellow Inc., #FP301200) with a 150 µm aperture was prepared. The aperture was pretreated by 0.5 µL, 10 % (v/v) hexadecane/pentane mixture and was dried immediately by N2. Then 500 µL, 10 mM Hepes buffer (pH 7.4, with 1 M KCl) and 5 uL DOPC:DOPG (4:1, 10 mg/mL) was added into the two compartments (*cis* and *trans*) of the home-made chamber. The single-channel electrical recording was carried out at room temperature through a pair of Ag/AgCl electrodes. The *cis* side was defined as the grounded side and a voltage potential was applied to the *trans* side, which means that the positively charged analytes translocate from *trans* to *cis* through the bilayer.

### 2.3 Single-channel electrical recordings and data analysis

Single-channel electrical recordings were conducted in a whole cell mode with a patch clamp amplifier (Axopatch 200 B, Axon instrument, Molecular Devices, CA). The purified recombinant full-length mPrP(23-230) or the fragmented PrP (FT(23-110), FT(23-50), FT(32-80), OR(53-90) or FT(93-110)) in the presence of 0.1 % DDM, was added to the *trans* part of the chamber with the final concentration 0.4 μM. 100 mV voltage was applied during all the experiments. For data acquisition, a DigiData 1440 A/D converter (Axon) was equipped with a PC, where the pClamp and Clampfit were installed for data process.

## 3 Results and discussion

### 3.1 PrP sequence with more amino acids remaining in the primary N-terminal domain shows less channel activity

To investigate the effect of different functional segments on PrP channel activity, single-channel electrical recordings were firstly carried out across physiologically relevant and negatively-charged membranes comprised of DOPC and DOPG at a ratio of 4:1 in 10 mM Hepes buffer (pH 7.4) with 1 M KCl (23). The two sides separated by lipid bilayer are *cis* or *trans* side, in which *cis* part is defined as the grounded side. The positively charged analytes translocate from *trans* to *cis* through the bilayer when a positive potential is applied. As full-length mPrP(23-230) and other functional fragments contain positive charges, we added the recombinant PrP into the *trans* part of the chamber where the channel formation was recorded with current trace under 100 mV. Abrupt jumps in the electrical recording were observed, attributed to channel formation, and the histograms of the counts against current were extracted unambiguously in **Fig. 2A**. mPrP(23-230) formed two similar channels with distinct conductance (0.02 and 0.04 nS), presumably caused by conformational transition of heterogeneous PrP in lipid bilayers (24). We next sought to investigate if the N terminus of prion protein (FT) was sufficient to induce channel activity. We tested full-length N terminus (FT(23-110)) or peptides spanning the entire FT region. With different functional segments of FT, we observed that full-length FT(23-110) (**Fig. 2B**) and FT(23-50) (**Fig. 2C**) sequences formed channels in the lipid bilayers, whereas other FT (residues 32-80, 53-90 or 93-120 in **Fig. 2D**) segments did not induce pores the lipid bilayers, indicative of the essential role of the proximal N-terminal amino acids (23-50) for PrP channel activity. The data support previous observations where the mice expressing the first nine amino acids (23–31) displays spontaneous neurodegenerative phenotypes (25), and the same segment resembles as a transduction domain to insert lipid membranes (11,12). In line with our studies, these studies support the role of FT (23–31) in PrP-related neurotoxicity. Among the mPrP segments and the full-length mPrP, the shortest fragment FT(23-50) formed the largest channel with a varied conductance of 2.25-13 nS, compared to that of FT(23-110) with 0.05-0.15 nS conductance. This reveals that FT(23-50) dominates the pore formation and the C-terminal domain or other FT sequence down-regulates the channel formation of FT(23-50) for the full-length channel activity. The stabilization of the GD domain or other FT region is therefore assumed to counteract the regulation, thereby enhancing the channel activity.

**Figure 2.**
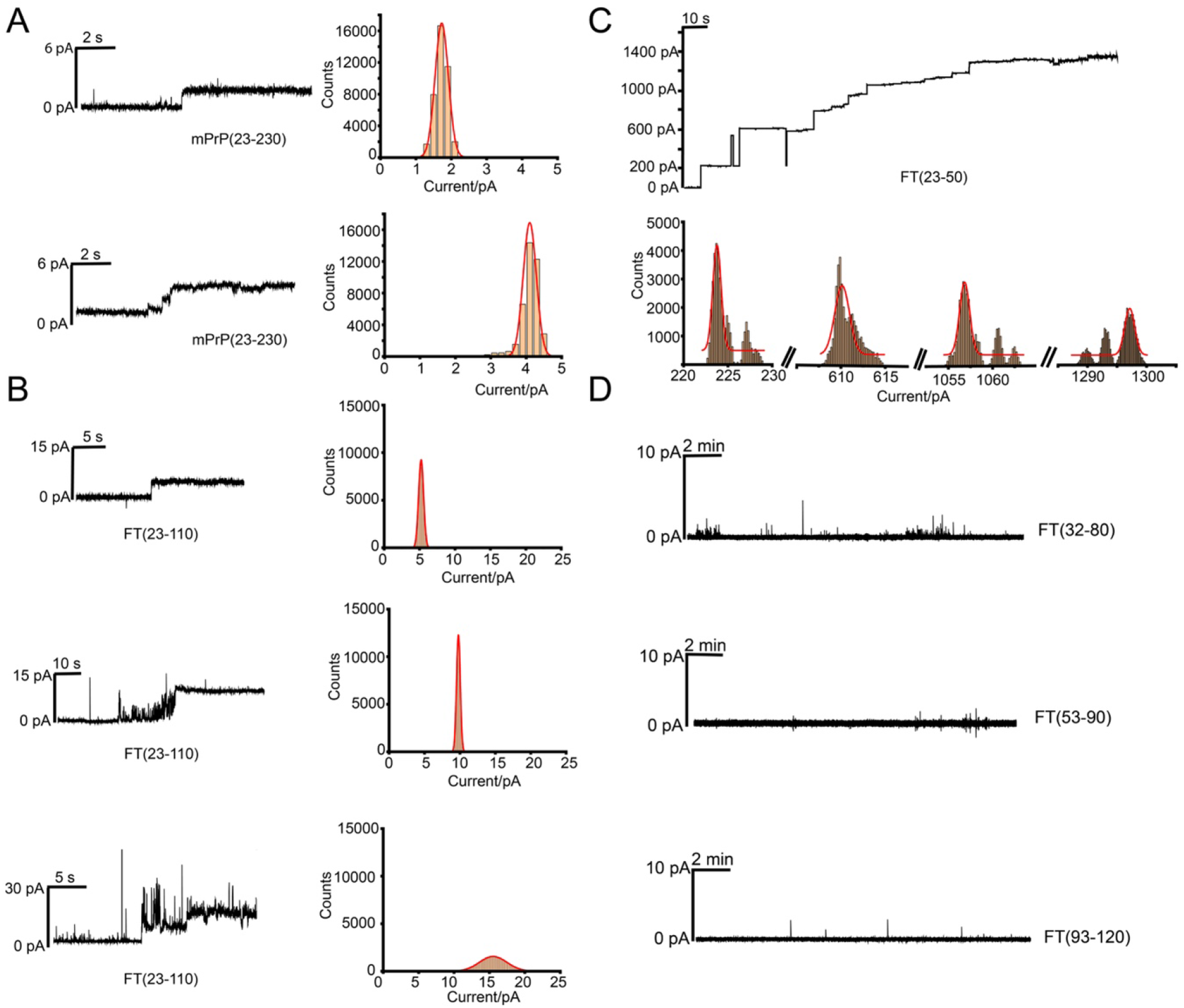
Representative current traces of single-channel electrical recordings induced by the full-length recombinant mPrP(23-230) protein (A) and the truncated N-terminal domains of PrP (B-D). Heterogeneous channels have been observed with subtle varied currents, indicative of different oligomeric states in the lipid membranes. Histograms of the counts generated by the induced current traces were also shown close to the current trace. The Gaussian function method was applied for fitting the distribution of these counts. The final concentration of the peptides in the *trans* side of the chamber is 0.4 µM in 10 mM Hepes buffer, pH 7.4 with 1 M KCl. Lipid bilayers were prepared by using a lipid mixture comprised of 10 mg/mL DOPC:DOPG at a ratio of 4:1.

### 3.2 C-terminal domain blocks the full-length mPrP(23-230) channel

To test this possibility, single-channel electrical recording was conducted for the full-length mPrP(23-230) in the presence of the GD region binding antibody, POM1 in **Fig. 2A**. This allowed determining if the full-length channel activity can be prompted through the stabilization of GD. Shown in **Fig. 3B**, we observed the significantly enhanced channel activity of mPrP(23-230) after the addition of POM1. This implies that POM1, upon binding to the globular domain, enhances the interaction of N-terminus interaction with lipid membranes. Overall, this result suggests that binding of POM1 induces a conformational change that might facilitate the interaction of N-terminus with the lipid bilayer and thereby induces pores. To further confirm this observation, the mutant antibody POM1_Y104A (26), which targets the same epitope with POM1 on mPrP(23-230) but is non-toxic, almost abolishes the channel activity of mPrP(23-230) in the lipid bilayers (**Fig**. 3D).This data suggests that POM1_Y104A does not induce the toxic conformation responsible for the interaction of the FT with lipid membranes. It suggests that the POM1_Y104A antibody induces the different conformation of GD segment, thereby C-terminus retains the activity to block the interaction between the FT segment and lipid membranes. Our study is in agreement with the previous work by Frontzek *et al*., who demonstrated the neuroprotective effect of POM1_Y104A on neuronal cells expressing the prion proteins (26). Briefly, we found that the antibodies regulate neurotoxicity of the prion proteins by stabilizing the GD domain and then further changing the FT(23-50) channel forming activity in lipid membranes.

**Figure 3.**
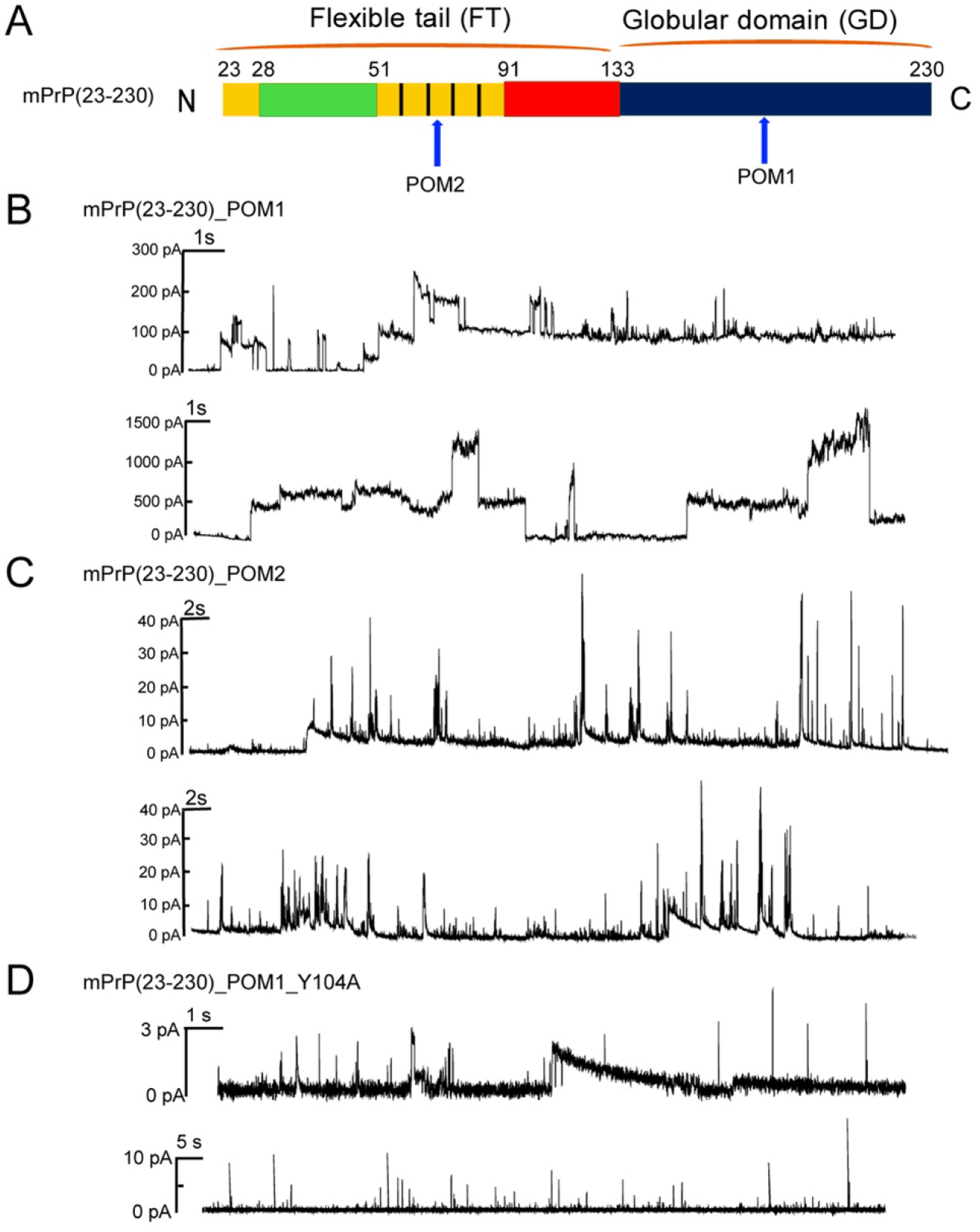
Schematic representative of single channel electrical recordings with the full-length recombinant mPrP(23-230) protein in the presence of antibodies. (A) Binding of POM1 and POM2 to GD and the octarepeats of FT, respectively. (B) Representative traces of spontaneous currents recorded from mPrP(23-230) with POM1 (B), POM2 (C) and POM1_Y104A. POM1 significantly enhances the mPrP(23-230) channel activity, POM2 gives negligible effect on channel formation and POM1_Y104A retains the channel activity. The proteins were added to the *trans* side up to a final concentration of 0.4 µM.

### 3.3 Octarepeats of PrP block the channel activity of FT (23-110) protein

We next investigated whether agents stabilizing FT can alter the FT(23-50) channel activity. To this end we used POM2 (binding to the OR region; residues 51-90), which was previously also shown to prevent neurodegeneration induced by prions and POM1. We found that unlike POM1, POM2 did not enhance the channel activity of mPrP(23-230) (**Fig. 3C**). This was out of our anticipation that the stabilization of OR domain by POM2 could increase mPrP(23-230) channel activity. But it is plausible that the binding of POM2 to OR only stabilize the FT domain but not the GD domain of mPrP(23-230), hindering the channel activity in the lipid bilayers. We next investigated how POM1 binding affects the pore formation by FT domain alone. FT(23-110) was used to determine the channel activity after the addition of the POM2 antibody in **Fig. 4A**. We found that FT(23-110) in the presence of POM2 antibody formed channel with a higher conductance (approximately 0.1-0.3 nS) (**Fig. 4B)** than that of mPrP(23-230) (at around 0.05 nS) in **Fig. 3C**. This confirms that the GD domain down-regulates the N-terminus channel formation for the full-length prion proteins in the lipid membranes. We therefore conclude that stabilizing the OR domain plays only a negligible role the channel activity of full-length mPrP(23-230) but clearly enhances the channel activity of the FT domain alone.

**Figure 4.**
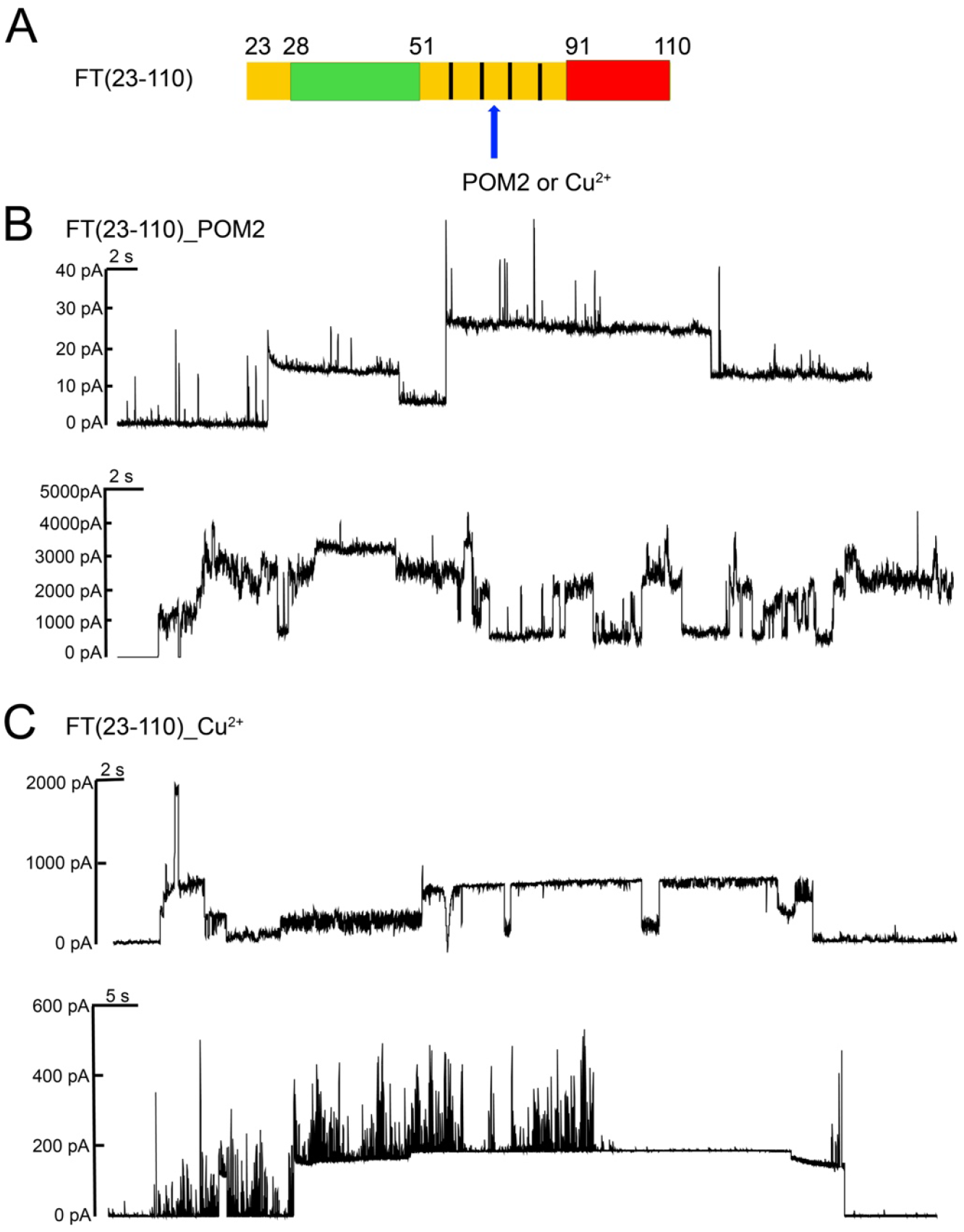
(A) Binding sites of POM2 and Cu^2+^ ions on the FT(23-110) sequence. (B-C) The representative traces of spontaneous currents recorded from mPrP(23-230) with POM2 (B) or with Cu^2+^ (C). POM2 or Cu^2+^ significantly enhances the FT(23-110) channel activity.

Besides POM2, Cu^2+^ ions bind to the OR region and regulate the PrP toxicity at physiological concentrations (27). We then tested if Cu^2+^ ions up- or down-regulates the channel activity of FT(23-110). Like POM2, Cu^2+^ ions up-regulated the channel activity of FT(23-110) in **Fig. 4C**. Our studies further support the conclusion that OR segment stabilization can enhance the neurotoxicity of FT(23-110) segment.

## 4 Discussion

Neurodegenerative disorders including prion diseases have been suggested to share, as a common neurotoxic mechanism, that the misfolded amyloid proteins interact with and permeabilize lipid membranes, leading to cell dysfunction (14). Several studies have shown that the primary N-terminal amino acids (23-31) are a prerequisite for channel activity (11,12), whilst the effect of GD and other FT region on the channel activity of the primary N-terminal region remains to be explored. By using single-channel electrical recordings in the presence of the different functional segments of FT, we investigated how FT and GD change the channel activity or membranes permeabilization of the primary region.

Our results show that the full-length recombinant mPrP(23-230) is less active than the N-terminal fragment FT(23-110). Additionally, the more amino acids added to FT(23-31), the less channel activity observed in lipid bilayer. FT(23-50) possesses the largest conductance (2.25-13 nS), indicative of the highest channel activity, and FT(23-110) or full-length mPrP(23-230) provokes lower conductance (down to 0.2 nS). This shows that GD or other FT sequence impairs the channel activity. Previously, Wu *et al*. reported (12) that truncated PrP(1-109) induces higher ionic currents on N2a cells than that of truncated PrP segment FT(1-90), followed by FT(1-58). We assume that the main discrepancy attributes to the artifact of EGFP (enhanced green fluorescence protein) molecule equipped with GPI (glycosylphosphatidylinositol) anchor linked to N-terminal PrP in the earlier study (12). Like other FT functional segments, the linked EGFP protein presumably modulate the interaction of the primary N-terminal amino acids FT(23-50) with lipid membranes. Our setup provides a more direct method without an external fused protein to study the channel activity of truncated PrP in the lipid membranes. The enhanced full-length mPrP(23-230) channel activity with C-terminal (POM1) rather than the N-terminal binding antibody (POM2) further supports the down-regulating effect of GD domain on the full-length mPrP channel activity in the lipid membranes. Moreover, binding of FT(23-110) to POM2 or Cu^2+^ increases the channel activity. This confirms the dominant role of GD on the N-terminus channel formation for the full-length prion proteins in the lipid membranes. The provoked spontaneous ionic currents agree with the previous observation where the toxic POM1 antibody acted as prion mimetic inducing neuronal death (28). This implies the underlying ion channel formation as the neurotoxic mechanism of the prion proteins. We also observed the small current conductance (0.02-0.04 nS) of full-length mPrP(23-230). These subtle changes could be negligible in a complicated environment of cellular studies (11,13). Our in-bulk study may provide a straightforward method for characterizing membranes permeabilization of the PrP proteins, revealing the molecular pathogenesis of prion disease as well as screening inhibitors against the disease.

In summary, our results reveal that the primary N-terminal FT(23-50) is essential for the membranes permeabilization. It is consistent with the determinant role of primary nine amino acids (23-31) on the channel activity (11). Second, the membranes permeabilization is negatively correlated with C-terminal extended functional segment of prion segment. Stabilization of the functional segment can be positively correlated with the membrane permeabilization, indicative of the molecular mechanism of prion neurotoxicity.

